# Assessing the impacts of various factors related to identification, conservation, biogenesis, and function on circular RNA reliability

**DOI:** 10.1101/2022.10.28.514164

**Authors:** Trees-Juen Chuang, Tai-Wei Chiang, Chia-Ying Chen

**Affiliations:** Genomics Research Center, Academia Sinica, Taipei 115201, Taiwan

**Keywords:** Circular RNA, Reliability, Non-co-linear junction, RNA treatment, Relative impacts

## Abstract

Circular RNAs (circRNAs) are non-polyadenylated RNAs with a continuous loop structure characterized by a non-co-linear back-splice junction (BSJ). While dozens of computational tools have been developed and identified millions of circRNA candidates in diverse species, it remains a major challenge for determining circRNA reliability due to various types of false positives. Here, we systematically assess the impacts of numerous factors related to identification, conservation, biogenesis, and function on circRNA reliability by comparisons of circRNA expression from mock (total RNAs) and the corresponding co-linear/polyadenylated RNA-depleted datasets based on three different RNA treatment approaches. Eight important indicators of circRNA reliability are determined. The relative contribution to variability explained analyses further reveal that the relative importance of these factors in affecting circRNA reliability is conservation level of circRNA > full-length circular sequences > supporting BSJ read count > both BSJ donor and acceptor splice sites at the same co-linear transcript isoforms > both BSJ donor and acceptor splice sites at the annotated exon boundaries > BSJs detected by multiple tools > supporting functional features > both BSJ donor and acceptor splice sites undergoing alternative splicing. By extracting RT-independent circRNAs, circRNAs passing multiple experimental validations, and database-specific circRNAs, we showed the additive effects of these important factors in determining circRNA reliability. This study thus provides a useful guideline and an important resource for selecting high-confidence circRNAs for further investigations.

## Introduction

CircRNAs are an emerging class of RNAs formed by *cis*-backsplicing with a covalently closed loop structure that are endogenously expressed as non-polyadenylated circular molecules (Chen et al. 2015; Chen 2020). Such a loop structure is characterized by a non-co-linear (NCL) back-splice junction (BSJ) between a downstream splice donor site and upstream splice acceptor site. Despite most circRNAs are expressed at a much lower level compared with linear mRNAs (Jeck et al. 2013; Salzman et al. 2013; Guo et al. 2014), some are highly expressed (Salzman et al. 2012; Jeck et al. 2013), even more abundant than their co-linear counterparts (Zheng et al. 2016; Chuang et al. 2018). In addition, circRNAs are more stable than their corresponding colinear mRNA isoforms (Chen et al. 2015; Chen and Yang 2015). Some circRNAs were further demonstrated to be evolutionary conserved across species (Jeck et al. 2013; Yu et al. 2014; Rybak-Wolf et al. 2015; Chen et al. 2020). Accumulating evidence demonstrated that circRNAs can participate in various aspects of biological processes, including in competing with their host gene expression and splicing, regulating activities of microRNAs (miRNAs) and RNA binding proteins (RBPs), and acting as translation templates (Chen 2020; Huang et al. 2020). CircRNAs were reported to be especially abundant in brain and neuronal tissues (Rybak-Wolf et al. 2015; You et al. 2015) and regulatory important in nervous system development and aging (Gasparini et al. 2020). Loss or dysregulation of circRNAs may affect brain function (Piwecka et al. 2017; Chen et al. 2020; Gasparini et al. 2020), indicating the roles of circRNAs in pathogenesis and progression of neurological diseases. In cancer studies, alternation of circRNA expression was shown to affect cell apoptosis, invasion, migration, and proliferation, revealing the potential of circRNAs for serving as biomarkers and therapeutic targets in cancer (Liu et al. 2020; Rajappa et al. 2020). These findings suggest that circRNAs are an ancient and fine-tuned class of functional transcripts.

Varying bioinformatics tools have been developed to identify BSJs using high-throughput transcriptome sequencing (RNA-seq) data (Chen et al. 2021), leading to a tremendous number of potential circRNA candidates in human and other species (Xia et al. 2018; Wu et al. 2020; Vromman et al. 2021). However, circRNA detections are hampered by various types of false positives. An NCL junction that originates from sequencing error, *in vitro* artifact, ambiguous alignment, *trans*-splicing, or genetic rearrangement is often misjudged as a BSJ event by computational detections (Chen et al. 2015; Chen 2020). This reflects the fact of limited overlap among the circRNA candidates identified by different tools (Chen et al. 2015; Hansen et al. 2016; Szabo and Salzman 2016; Zeng et al. 2017; Chen and Chuang 2019a) or databases (Vromman et al. 2021), highlighting the issue of circRNA reliability. Although millions of circRNA candidates were deposited in varied databases, only a very limited number of candidates have been carefully curated to date (Vromman et al. 2021). It requires a guideline for evaluating the reliability of the detected circRNA candidates before performing time- and cost-consuming experimental validations.

Since circRNAs are NCL RNAs and lack 3’polyadenylated tails, many circRNA candidates were detected in RNA-seq datasets derived from total RNAs (the ribosomal RNA-depleted RNAs without poly(A)-selection; “mock RNAs” for simplicity), which contain RNA-seq reads from both co-linear and NCL RNAs (Memczak et al. 2013). Because most co-linear RNAs are digested by RNase R (Jeck et al. 2013; Memczak et al. 2013; Zhang et al. 2014; Zhang et al. 2016a), a detected circRNA candidate is more likely to be real if its normalized BSJ read count is enriched or not reduced after the treatment. Therefore, comparisons of BSJ read counts detected from mock and RNase R-treated RNA datasets were often employed to evaluate the reliability of the detected circRNAs (Hansen et al. 2016; Zeng et al. 2017; Chen and Chuang 2019a; Jakobi et al. 2019). However, in some cases such a mock-treated comparison approach may not be efficient for circRNA detection due to two inherent limitations. First, some co-linear RNAs containing highly structured 3’ ends or G-quadruplexes are also resistant to RNase R (Xiao and Wilusz 2019). Second, some circRNAs may be susceptible to RNase R and digested by this exonuclease (Zhang et al. 2016b). Accordingly, in addition to RNase R-treated RNAs, we considered two other RNA treatments, RNAs treated with coupling A-Tailing and RNase R digestion and RNAs treated with depletion of both ribosomal RNAs and polyadenylated RNAs, for mock-treated comparisons. The former could efficiently digest most the RNase R-resistant co-linear RNAs stated above (Xiao and Wilusz 2019); and the latter could deplete most polyadenylated RNAs and retained circRNAs susceptible to RNase R (Zhang et al. 2016b). On the basis of comparisons of normalized BSJ read counts detected from mock and the corresponding co-linear/poly(A) depleted RNA datasets from three different RNA treatments, we thus presented a systematical workflow to evaluate the impacts of various factors related to circRNA identification, conservation, biogenesis, and function on circRNA reliability. After determining statistically-important indicators of circRNA reliability, we further assessed the relative influence of these important indicators on circRNA reliability. Collectively, we provided a useful guideline for selecting high-confidence circRNAs. All of the information related to circRNA identification, conservation, biogenesis, and function was freely available.

## Results

### Description of the proposed workflow and used datasets

As shown in our analysis flow (Fig. 1A), we first extracted circRNA (or BSJ) candidates from the circAtlas database 2.0 (Wu et al. 2020) (see also Supplemental Table S1), which contained 580,654 human circRNA candidates based on 240 samples collected from 19 tissues across seven vertebrate species. CircAtlas circRNA candidates were identified by four bioinformatics tools including CIRI2 (Gao et al. 2015; Gao et al. 2018), find_circ (Memczak et al. 2013), circExplorer (Dong et al. 2019), and DCC (Cheng et al. 2016). Like previous reports (Chen et al. 2015; Hansen et al. 2016; Szabo and Salzman 2016; Zeng et al. 2017; Chen and Chuang 2019a; Rabin et al. 2021), the majority (63.5%) of the examined circRNA candidates were detected by only one or two tools (Fig. 1B). Such great discrepancies in the identification results among tools reflected the uncertainty of the computationally detected circRNA events (Chen et al. 2015; Hansen et al. 2016; Szabo and Salzman 2016; Zeng et al. 2017; Chen and Chuang 2019a). One cause of the false positive calls may be due to alignment ambiguity with an alternative co-linear explanation or multiple hits when detecting BSJs (Chen et al. 2015; Chen and Chuang 2019b; Chen and Chuang 2019a). Regarding the 580,654 circAtlas circRNAs, we found that 17.3% (100,183 circRNAs; Fig. 1C) were potential false positives arising from an alternative co-linear explanation or multiple hits (Methods). It was reported that the circRNAs detected by multiple tools tended to be more reliable than tool-specific circRNAs (Petkovic and Muller 2015; Hansen 2018). As expected, the percentages of circRNAs derived from ambiguous alignments significantly decreased with increasing the numbers of the detected tools (Fig. 1D), reflecting the positive correlation between the level of circRNA reliability and the number of circRNA detection tools. For accuracy, we excluded the 100,183 circRNAs and considered the remaining circRNAs (480,471 events) for the following analyses.

**Figure 1.**
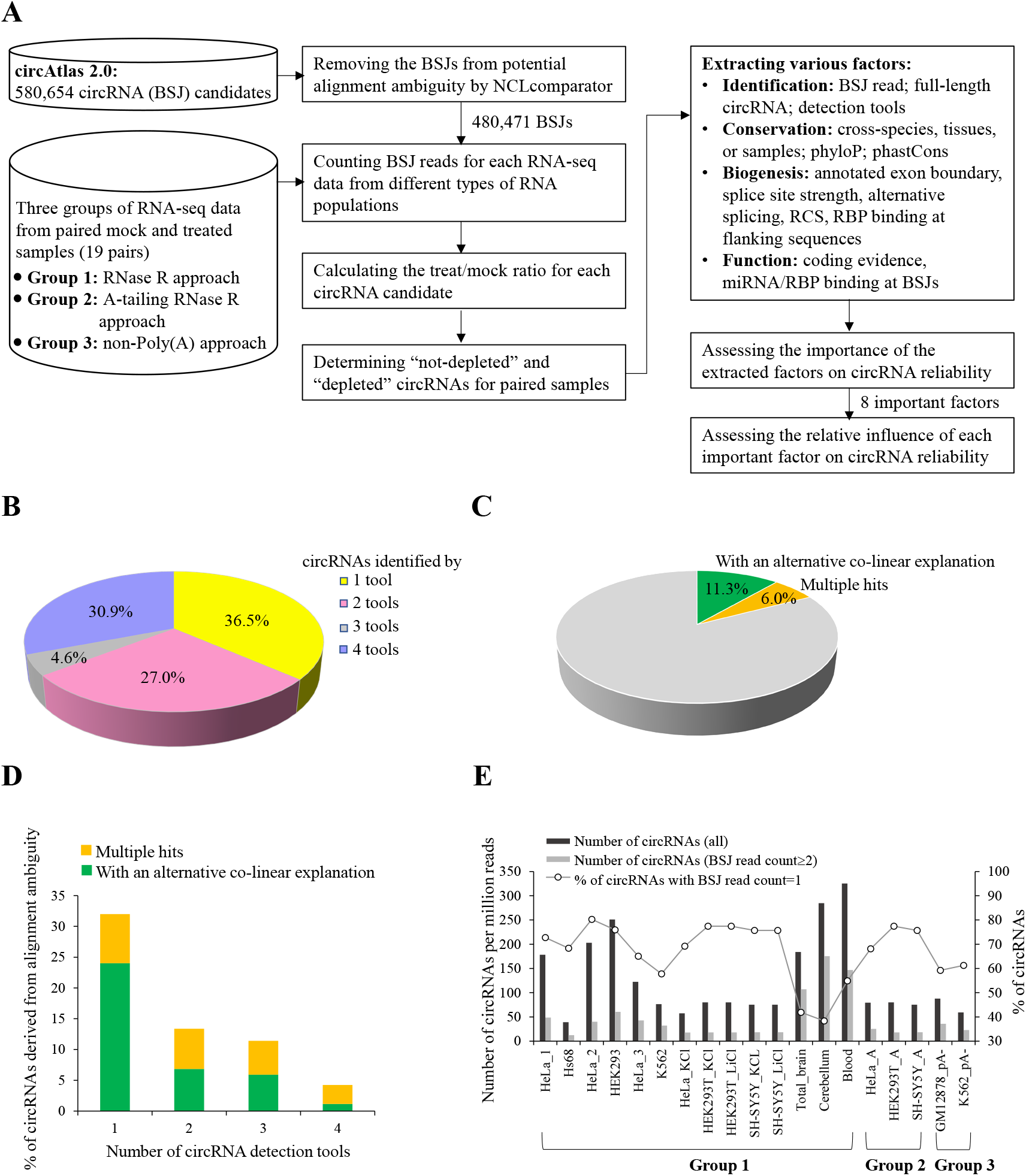
The assessment of the impacts of various features on circRNA reliability. (A) Flowchart of the overall analyses. (B) The distribution of the extracted circAtlas circRNA (BSJ) candidates (580,654 candidates) identified by 1, 2, 3, or 4 circRNA detection tools. (C) The 580,654 candidates derived from potential alignment ambiguity (with an alternative co-linear explanation or multiple hits). (D) Comparisons of the percentages of circRNA candidates derived from potential alignment ambiguity for the candidates detected by 1, 2, 3, or 4 tools. (E) Comparisons of normalized numbers of circRNA candidates with supporting BSJ read count =1 or ≥ 2 in all extracted mock samples of the 19 mock-treated sample pairs. A considerable percentage of circRNA candidates (38%~80%) were supported by only one BSJ read.

Most circRNA candidates were detected from RNA-seq datasets derived from mock samples (the ribosomal RNA-depleted RNAs without poly(A)-selection), which contain RNA-seq reads from both circRNAs and co-linear/polyadenylated RNAs (Memczak et al. 2013). We employed comparisons of normalized BSJ read counts detected from mock and the corresponding co-linear/poly(A) depleted datasets to determine high-confidence circRNAs. For comprehensively assessing circRNA reliability, we extracted 19 mock-treated sample pairs of different samples or studies from public datasets and categorized them into three groups according to the RNA treatment approaches (see Table 1 and Supplemental Table S2):

1. **Group 1:** RNase R approach, mock sample vs. RNase R-treated sample (14 pairs).
2. **Group 2:** A-tailing RNase R approach, mock sample vs. A-tailing RNase R-treated (RNAs treated with coupling A-Tailing and RNase R digestion) sample (3 pairs).
3. **Group 3:** non-Poly(A) approach, mock sample vs. non-Poly(A)-selected (RNAs treated with depletion of both ribosomal RNAs and polyadenylated RNAs) sample (2 pairs).

**Table 1.**
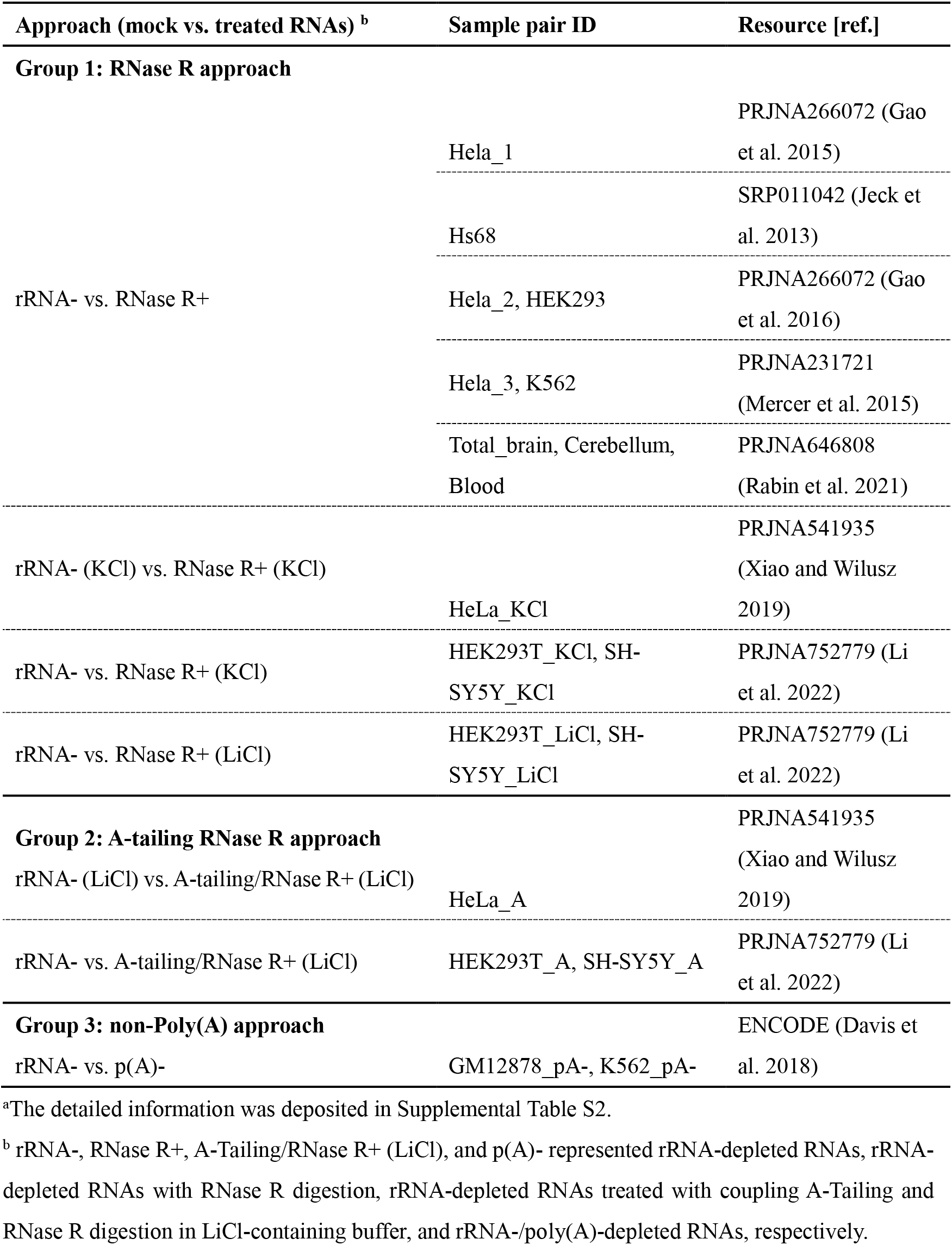
Summary of RNA-seq data used in this study ^a^.

We examined the 480,471 circRNAs in all the mock samples (Methods) and found that a considerable percentage of circRNAs (38%~80%) were supported by only one BSJ read (Fig. 1E). To minimize potential false positives, only the circRNAs expressed in the mock samples with the supporting BSJ read count ≥ 2 were considered in the analysis for each mock-treated sample pair. For each considered circRNA candidate, we calculated the ratio of the circRNA expression (the BSJ reads per million raw reads; RPM) detected in the treated samples to that detected in the corresponding mock samples (i.e., treat/mock ratio). It was presumed that the circRNAs not depleted after the treatment (“not-depleted circRNAs” or treat/mock ratio ≥1) were more likely to be *bona fide* circRNAs than those depleted due to the treatment (“depleted circRNAs”; or treat/mock ratio <1). We then compared not-depleted circRNAs with depleted ones in terms of various factors related to circRNA identification, conservation, biogenesis, and function (Supplemental Table S3), respectively; and thereby accessed the importance of these factors in affecting circRNA reliability.

### Factors related to circRNA identification

Regarding the circRNAs identified in the mock samples, the impacts of the factors related to circRNA identification (i.e., BSJ read count, number of detected tools, and evidence of full-length circular sequence) on circRNA reliability were assessed. First, we found that the supporting BSJ read counts were significantly higher in not-depleted circRNA candidates than in depleted ones in all mock-treated sample pairs examined (all FDR < 0.05; Fig. 2A). The percentages of non-depleted circRNAs increased with increasing the supporting BSJ read counts (Fig. 2A). Second, we examined the influences of BSJs detected by multiple or single circRNA detection tools on circRNA reliability and found that the former were significantly enriched for not-depleted circRNAs (Fig. 2B). Generally, the percentages of non-depleted circRNAs increased with increasing numbers of the detected tools (Fig. 2B). Third, we examined the influences of BSJs with or without the evidence of full-length circular sequences reconstructed by CIRI-full (Zheng et al. 2019) on circRNA reliability and showed that the former were significantly enriched for not-depleted circRNAs, regardless of the RNA treatment approaches (Fig. 2C). Since the performance of CIRI-full, a short read-based approach, for identifying full-length sequences of circRNAs is hampered by the length of short reads (Zheng et al. 2019), we also extracted the full-length circRNA candidates identified by circFL-seq based on nanopore long reads (Liu et al. 2021). Such a long read-based approach provided the direct support of full-length circular sequences from single-molecule long reads. Similarly, the BSJ events with the evidence of full-length circular sequences showed a significant enrichment for not-depleted circRNAs in all mock-treated sample pairs examined (Fig. 2C).

**Figure 2.**
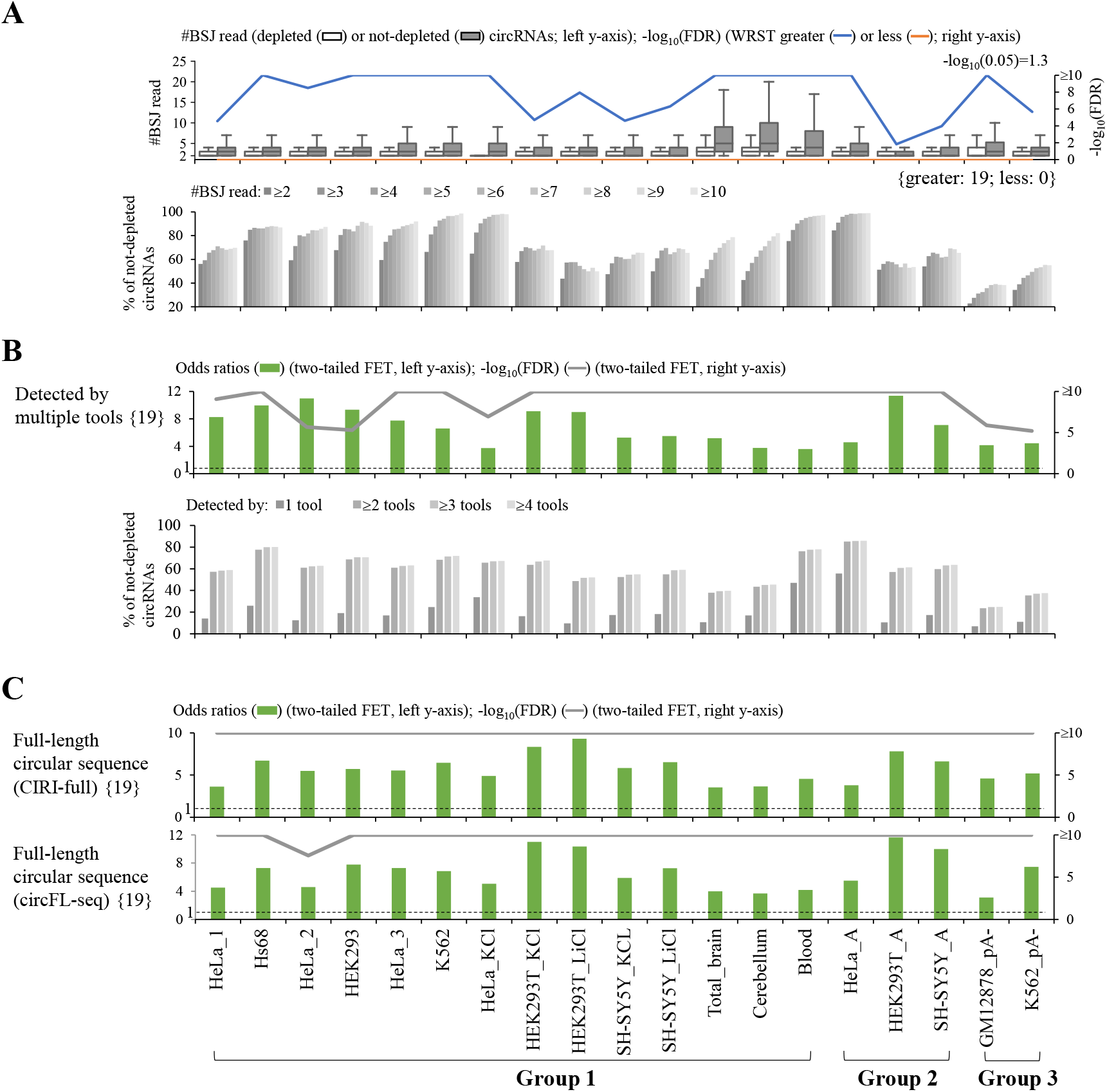
Impact of factors related to circRNA identification on circRNA reliability: (A) supporting BSJ read count, (B) number of detected tools, and (C) evidence of full-length circular sequence. For (B) and (C), the odds ratios, which were determined using two-tailed Fisher’s exact test (FET), represented the ratios of the occurrences of (B) circRNAs detected by multiple tools or (C) circRNAs with the evidence of full-length circular sequence for non-depleted circRNAs to the occurrences of those for depleted circRNAs. The dashed lines represented odds ratio=1. For the bottom panels of (A) and (B), the correlations between percentages of non-depleted circRNAs and (A) supporting BSJ read count or (B) number of detected tools were shown. For (C), the evidence of full-length circular sequence was supported by CIRI-full (a short read-based approach, top) or circFL-seq (a long read-based approach, bottom). *P* values were determined using Wilcoxon rank-sum test (WRST; greater or less; (A)) or two-tailed FET ((B) and (C)) and FDR adjusted across 19 mock-treated sample pairs for each examined factor using Benjamini-Hochberg correction. The number of mock-treated sample pairs that passed the statistical significance tests with FDR<0.05 were represented in curly brackets.

### Factors related to circRNA conservation

We then examined the relationship between circRNA reliability and conservation patterns at the three levels: species, tissues, and individuals (or samples). We found that the conservation levels of circRNAs at the three levels (Figs. 3A–3D) were all significantly higher in not-depleted circRNAs than in depleted ones, regardless of the RNA treatment approaches. The percentages of non-depleted circRNAs increased with increasing numbers of the conserved species (Fig. 3A), tissues (Fig. 3B), and samples (Fig. 3D). Of note, in contrast with the number of tissues (Fig. 3B), the levels of tissue specificity index were significantly lower in not-depleted circRNA than in depleted ones (Fig. 3C). We further showed that the evolutionary rates (as determined by phyloP (Pertea et al. 2011) or phastCons (Siepel et al. 2005)) of the sequences around the BSJs (see Methods) were generally higher in not-depleted circRNAs than in depleted ones (Supplemental Fig. S1). These results suggested that the BSJ events widely detected across species/samples were more reliable than those detected in a limited number of species/samples.

**Figure 3.**
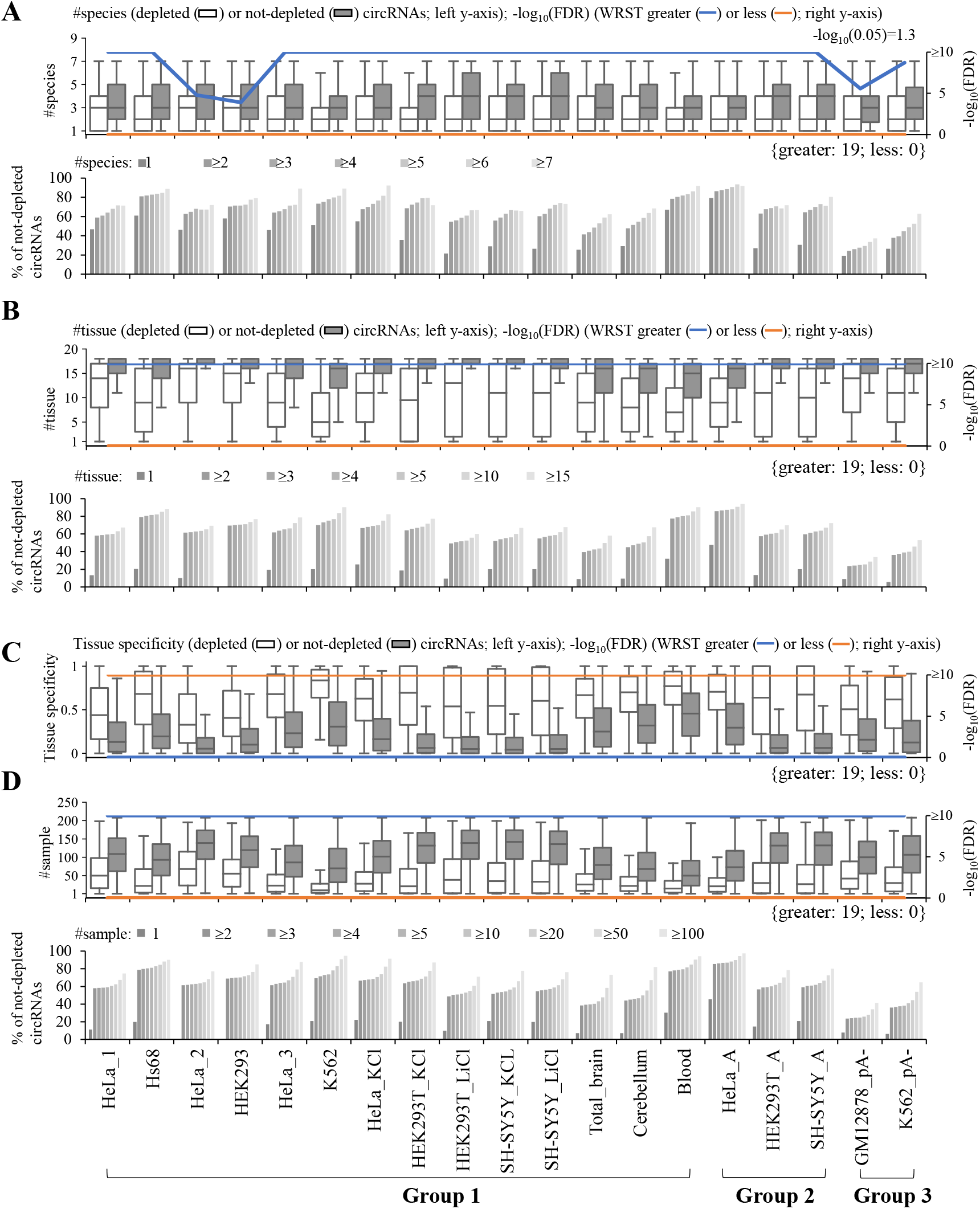
Impact of conservation patterns at the (A) species, (B and C) tissue, and (D) individual (or sample) levels on circRNA reliability. For the bottom panels of (A), (B), and (D), the correlations between percentages of non-depleted circRNAs and (A) number of conserved species, (B) number of conserved tissues, or (C) number of conserved samples were shown. *P* values were determined using WRST (greater or less) and FDR adjusted across 19 mock-treated sample pairs for each examined factor using Benjamini-Hochberg correction. The number of mock-treated sample pairs that passed the statistical significance tests with FDR<0.05 were represented in curly brackets.

### Factors related to circRNA biogenesis

We proceeded to examine the relationships between circRNA reliability and the factors related to circRNA biogenesis. Since circRNAs were suggested to be generated by canonical spliceosomal machinery (Ashwal-Fluss et al. 2014; Chen et al. 2015; Starke et al. 2015; Wang and Wang 2015), we examined whether the BSJs of not-depleted circRNA candidates tended to agree to well-annotated exon boundaries of co-linear transcripts or be subject to alternative splicing (AS) based on previously annotated co-linear transcripts (e.g., the Ensembl annotation) as compared with those of depleted circRNA ones. Indeed, the BSJs that agreed to annotated exon boundaries (either donor or acceptor splice sites) showed a significant enrichment for not-depleted circRNAs, regardless of the RNA treatment approaches (Fig. 4A and Supplemental Fig. S2A). The BSJs with both donor and acceptor splice sites matching annotated exon boundaries from the same co-linear transcript isoforms were also enriched (Fig. 4B). We also observed that the splice site strength of BSJs was generally stronger in not-depleted circRNAs than in depleted ones, regardless of the tools used for estimating the splice site strength (Supplemental Fig. S2B). Regarding the BSJs matching previously annotated exon boundaries of co-linear transcripts, the BSJs (donor or acceptor splice sites) that were subject to AS were significantly enriched for not-depleted circRNAs across the mock-treated sample pairs examined (Supplemental Fig. S2C). Such a trend exhibited particularly enriched when both donor and acceptor splice sites of the BSJs were subject to AS (Fig. 4C). These observations support that most circularized exons are recognized by the spliceosome during canonical splicing.

**Figure 4.**
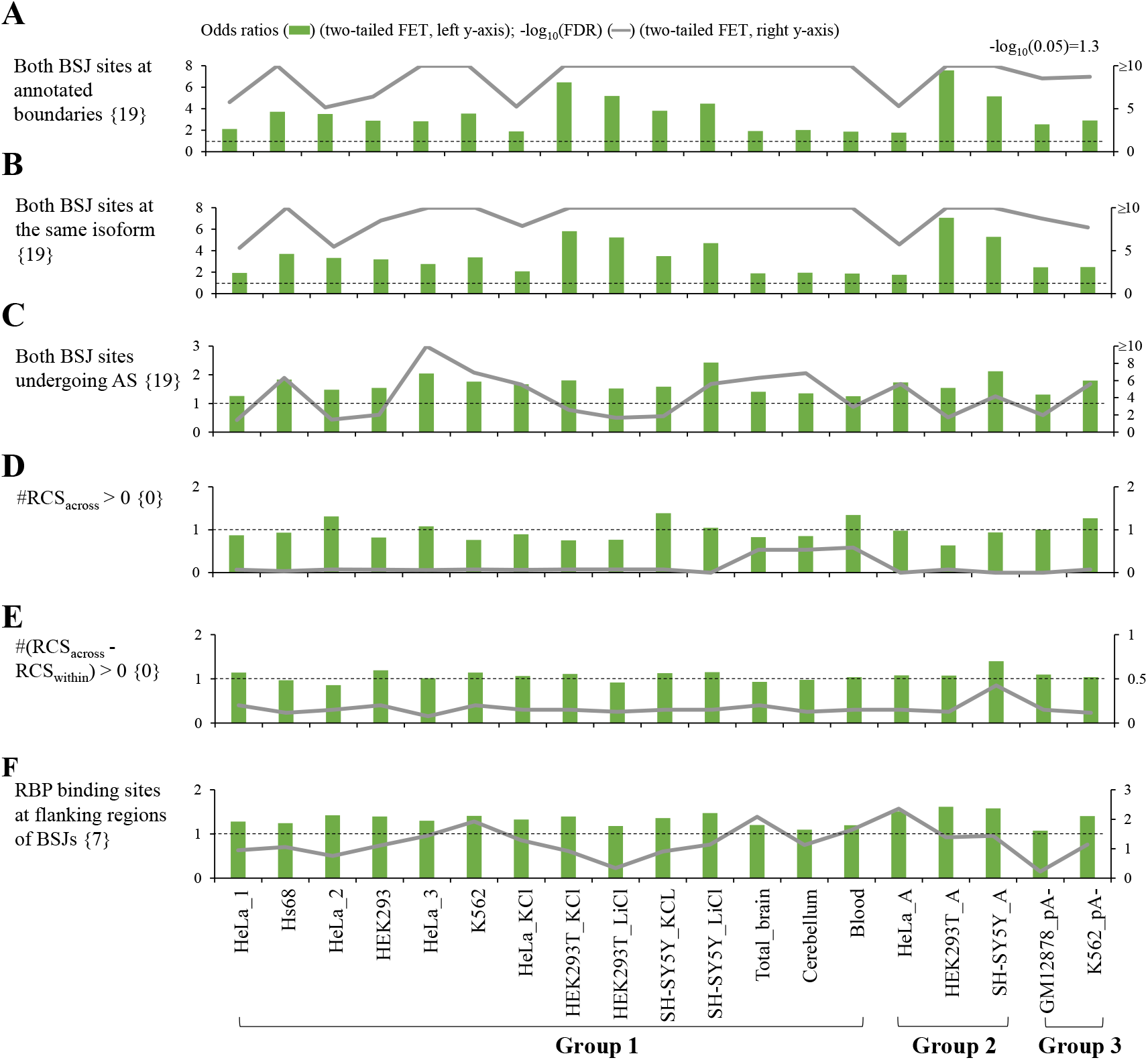
Impact of factors related to circRNA biogenesis on circRNA reliability. The odds ratios represented the ratios of the occurrences of (A) both BSJ donor and acceptor splice sites that agreed to well-annotated exon boundaries of co-linear transcripts, (B) both BSJ donor and acceptor splice sites that were derived from the same co-linear transcript isoforms, (C) both BSJ donor and acceptor splice sites that were subject to alternative splicing (AS), (D) BSJs that satisfied #RCS_across_>0, (E) BSJs that satisfied #(RCS_across_ - RCS_within_)>0, and (F) BSJs that had RBPs binding to the flanking regions for non-depleted circRNAs to the occurrences of those for depleted circRNAs, respectively. The dashed lines represented odds ratio=1. Odds ratios and *P* values were determined using two-tailed FET. *P* values were FDR adjusted across 19 mock-treated sample pairs for each examined factor using Benjamini-Hochberg correction. The number of mock-treated sample pairs that passed the statistical significance tests with FDR<0.05 were represented in curly brackets.

Next, previous studies demonstrated that back-splicing can be facilitated by RCSs residing in the sequences flanking circularized exons (Jeck et al. 2013; Zhang et al. 2014; Chuang et al. 2018) and affected by the competition of RCSs across flanking regions (RCS_across_) or within individual regions (RCS_swithin_) (Zhang et al. 2014). We thus asked whether the BSJs with RCS_across_ in flanking regions were enriched for not-depleted circRNA candidates. However, we did not observe this trend (Fig. 4D). We further examined the numbers of (RCS_across_ - RCS_swithin_) for the detected circRNAs and found no significant differences between not-depleted and depleted circRNAs in terms of the percentages of the BSJs with (RCS_across_ - RCS_swithin_) ≥ 1 (Fig. 4E). In addition, a number of RBPs were demonstrated to serve as a role for regulating circRNA biogenesis by binding to *cis* elements in the sequences flanking circularized exons (Chen et al. 2015; Chen 2020). We then examined whether the feature of BSJs with RBPs binding to the flanking regions showed an enrichment for not-depleted circRNAs and found that this feature was slightly enriched for not-depleted circRNAs only (Fig. 4F). These results suggested that the features of RCS and RBPs binding to the flanking regions were not good indicators for evaluating circRNA reliability.

### Factors related to circRNA function

We then examined the impact of functional features (miRNA binding, RBP binding, and seven types of evidence for circRNA coding potential) on circRNA reliability (Methods). For each BSJ event, since the circularized sequence is shared with its co-linear counterpart, we considered the predicted miRNA/RBP binding sites and ORFs spanning the BSJ. Integration of these nine types of functional features, we observed that numbers of supporting functional features were significantly higher in not-depleted circRNAs than in depleted ones in all mock-treated sample pairs (all FDR < 0.05; Fig. 5). The percentages of non-depleted circRNAs increased with increasing numbers of supporting functional features (Fig. 5). These results suggested that the strength of functional evidence for circRNA candidates were positively correlated with the level of circRNA reliability.

**Figure 5.**
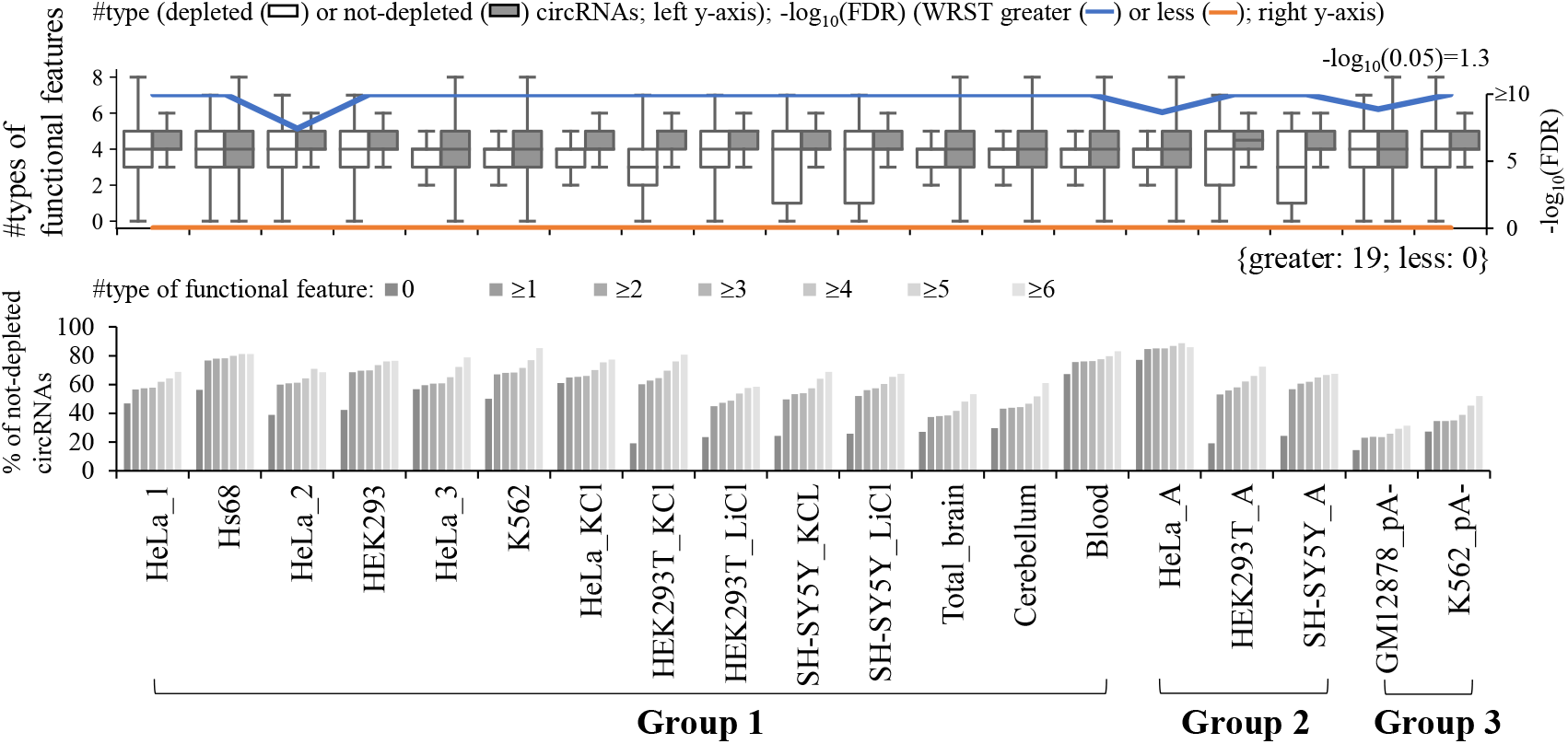
Impact of functional features on circRNA reliability. Nine types of functional features (see Methods) for circRNAs were examined. For the bottom panel, the correlations between percentages of non-depleted circRNAs and number of supporting functional features were shown. *P* values were determined using WRST (greater or less) and FDR adjusted across 19 mock-treated sample pairs for each examined feature using Benjamini-Hochberg correction. The number of mock-treated sample pairs that passed the statistical significance tests with FDR<0.05 were represented in curly brackets.

### Relative effect of each individual factor on circRNA reliability

According to the above assessment results, we observed eight important indicators of circRNA reliability, each of which can effectively distinguish between non-depleted and depleted circRNAs in all mock-treated sample pairs examined. The eight factors were (i) supporting BSJ read count; (ii) BSJs detected by multiple tools; (iii) full-length circular sequences; (iv) conservation level of circRNA (number of samples observed the circRNAs); (v) both BSJ donor and acceptor splice sites at the annotated exon boundaries; (vi) both BSJ donor and acceptor splice sites at the same co-linear transcript isoforms; (vii) both BSJ donor and acceptor splice sites undergoing annotated AS events; and (viii) supporting functional features. We then employed a generalized linear model (GLM) with the eight factors and measured the relative contributions to variability explained (RCVE) (Kvikstad et al. 2007) to evaluate the relative effect of individual factor in affecting circRNA reliability for each mock-treated sample pair (Methods) (Fig. 6A and Supplemental Table S4). We ranked the RCVE values for each mock-treated sample pair and found that on average the most important features in descending order were conservation level of circRNA, full-length circular sequences, supporting BSJ read count, both BSJ donor and acceptor splice sites at the same co-linear transcript isoforms, supporting functional features, both BSJ donor and acceptor splice sites at the annotated exon boundaries, BSJs detected by multiple tools, and both BSJ donor and acceptor splice sites undergoing annotated AS events (Fig. 6B).

**Figure 6.**
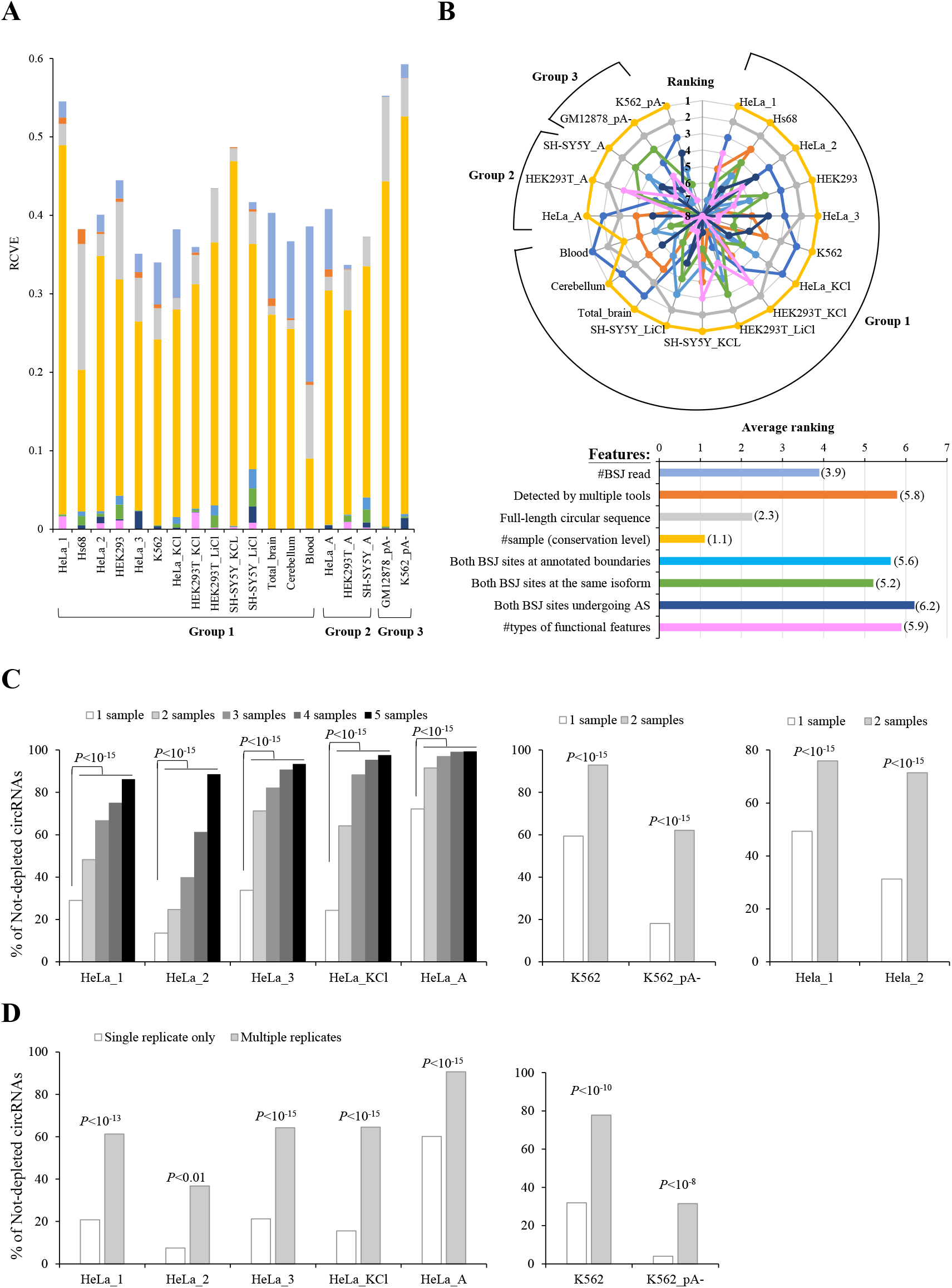
Assessment of relative influence of each individual factor on circRNA reliability. (A) The RCVE scores of the examined factors for each mock-treated sample pair. (B) The ranking (top) and average ranking (bottom; see also the numbers in the parentheses) of the RCVE scores of the examined factors. (C) The correlations between the percentages of not-depleted circRNAs and number of samples observed the circRNAs in Hela-based (left and right) or K562-based (middle) mock-treated sample pairs. Of note, the Hela_1 and Hela_2 mock-treated sample pairs were generated from the same group but different studies. (D) Comparisons of percentages of not-depleted circRNAs for Hela-specific circRNA (left) or K562-specific circRNAs (right) detected in single replicate only or multiple replicates. All *P* values were determined using two-tailed FET.

Particularly, our result revealed that conservation level of circRNA (number of samples observed the circRNAs) is the most dominant determinant of circRNA reliability common to all the examined mock-treated sample pairs except for the sample pair in blood (Fig. 6B). Of note, the samples considered were the integration of samples across species, tissue, and individuals in circAtlas. It is known that circRNAs are often expressed in a cell type-/tissue-specific manner (Salzman et al. 2013; Xia et al. 2017). Actually, we found that the majority (61%; 293,152 circRNAs) of the 480,471 circAtlas circRNAs were detected in one sample only (Supplemental Table S3). We were curious about whether the positive correlation between the percentage of not-depleted circRNAs and the number of samples observed the circRNAs held when considering one cell type/tissue only. Of the examined mock-treated sample pairs, the RNA-seq data of five pairs (Hela_1, Hela_2, Hela_3, Hela_KCl, and Hela_A) were all derived from Hela cells, which were generated from different studies or conditions (Supplemental Table S2). The five Hela-based RNA-seq datasets can be regarded as replicates. As expected, the percentage of not-depleted circRNAs was strongly positively correlated with the number of samples observed the circRNAs in all the five Hela-based pairs (Fig. 6C, left). The similar trend was observed for the two K562-based pairs (Fig. 6C, middle). We also examined the Hela_1 and Hela_2 mock-treated sample pairs, which were two replicates generated from the same group but different studies (Supplemental Table S2), and showed the similar results (Fig. 6C, right). Furthermore, regarding the mock samples of the 19 mock-treated sample pairs, we extracted the Hela-specific circRNAs that were detected in the examined Hela samples only but not in the non-Hela cell lines/tissues. For each of the five Hela-based mock-treated sample pairs, the Hela-specific circRNAs were divided into circRNAs detected in single replicate only and circRNAs detected in multiple replicates. Our results revealed that the percentages of not-depleted circRNAs were significantly higher in circRNAs detected in multiple Hela replicates than in circRNAs detected in single replicate only for all five Hela-based pairs (Fig. 6D, left). Such a trend held for the K562-based pairs (Fig. 6D, right).

## Discussion

The reliability of computationally detected circRNAs is not guaranteed due to varied types of false positives. It is particularly difficult to pick out spurious BSJ (or circRNA) events derived from *in vitro* artifacts among a tremendous number of previously identified NCL junctions. Such *in vitro* artifacts may frequently arise from template switching by random hexamer primed reverse transcriptase (RT) reactions (Houseley and Tollervey 2010), which can switch from one template to another with short homologous sequences at the NCL junction sites (McManus et al. 2010; Wu et al. 2014). Of note, some NCL junctions passing RT-based validations were finally confirmed to be generated from *in vitro* artifacts by further validations (Yu et al. 2014). As RNA-seq data are generated from RT-based sequencing strategies, it is unavoidable that a number of spurious NCL events masquerade as genuine BSJs due to such RT-based artifacts (Chen et al. 2015). We asked whether the approaches based on comparisons of paired mock and treated samples can reflect circRNA reliability to a certain extent. To this end, we extracted BSJs from two high-confidence circRNA datasets, in which the BSJ events had passed multiple experimental validations. For the first dataset (Chuang et al. 2018), BSJs were supported by both Avian Myeloblastosis Virus (AMV)- and Moloney Murine Leukemia Virus (MMLV)-derived RNA-seq reads (designated as “RT-independent circRNAs”). RT-independent circRNAs are regarded to be highly accurate because comparisons of different RTase products can effectively detect RT-based artifacts (Houseley and Tollervey 2010; Wu et al. 2014; Yu et al. 2014; Chen et al. 2015). For the second dataset (Fan et al. 2021), BSJs (or their functions) were previously validated by RT-based experiments and at least one type of non-RT-based experiments simultaneously in at least one human tissues or cell lines (designated as “RT-/non-RT-validated circRNAs”; Methods). We observed that the percentages of both RT-independent (Fig. 7A) and RT-/non-RT-validated (Fig. 7B) circRNAs were significantly higher in not-depleted circRNAs than in depleted circRNAs in all mock-treated sample pairs examined. Moreover, we compared circAtlas circRNAs with circRNAs from eight other publicly-accessible circRNA databases and identified circAtlas-specific circRNAs. We presumed that circAtlas-specific circRNAs were less probably to be reliable as compared with circRNAs collected in multiple databases. Indeed, the percentages of CircAtlas-specific circRNAs were significantly lower in not-depleted circRNAs than in depleted circRNAs (Fig. 7C).

**Figure 7.**
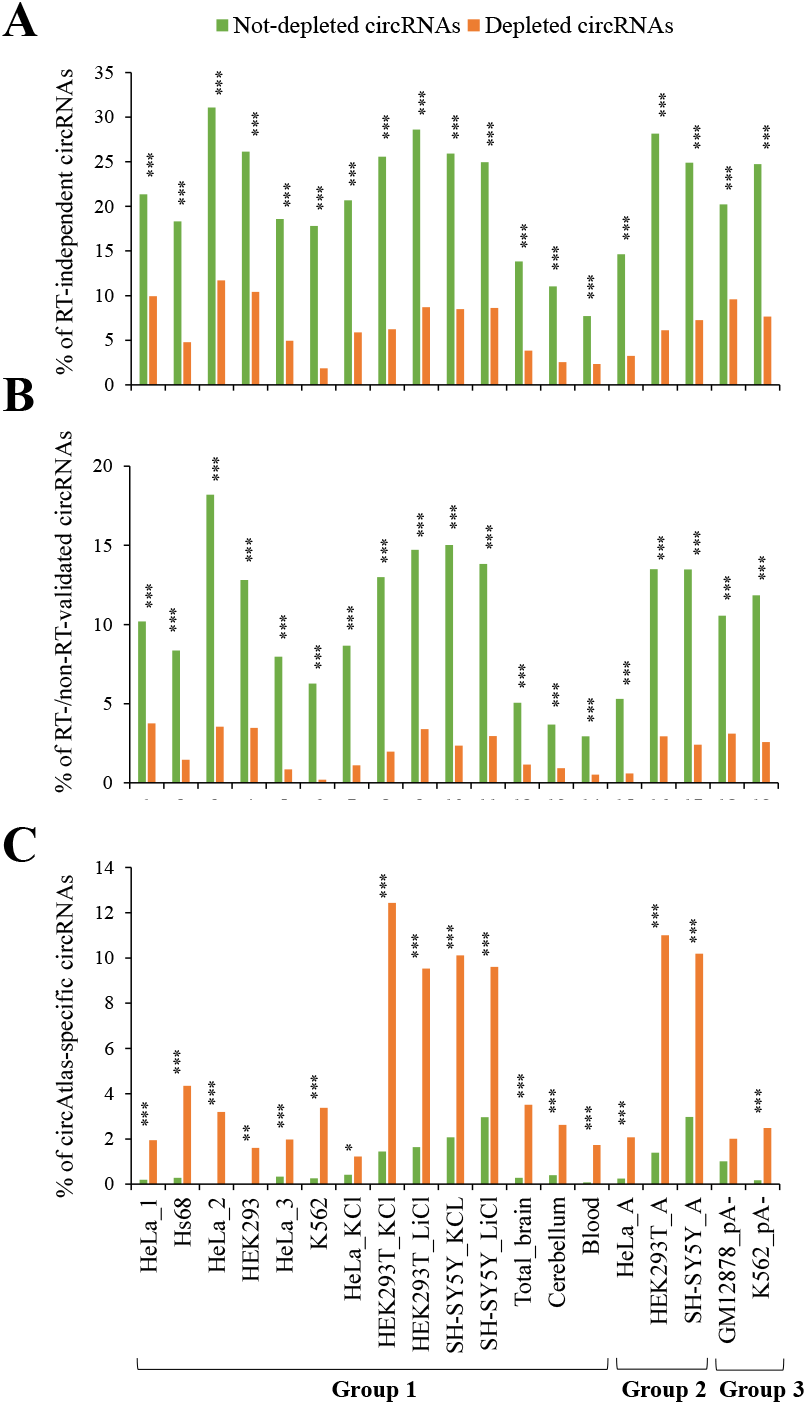
Comparisons of the percentages of (A) RT-independent circRNAs, (B) RT-/non-RT-validated circRNAs, and (C) circAtlas-specific circRNAs in the not-depleted and depleted circRNAs for all mock-treated sample pairs examined. RT-independent and RT-/non-RT-validated circRNAs represented high-confidence circRNAs (see text and Methods). CircAtlas-specific circRNAs were the circAtlas circRNAs that are not observed in eight other publicly-accessible circRNA databases (Methods). *P* values were determined using two-tailed FET and FDR adjusted across 19 mock-treated sample pairs for each examined feature using Benjamini-Hochberg correction. *, FDR<0.05. **, FDR<0.01. ***, FDR<0.001.

For the relative influence of each individual factor on circRNA reliability, our RCVE analysis revealed that conservation level of circRNA and full-length circular sequences were the top two dominant determinants of circRNA reliability (Fig. 6B). For the most important factor in affecting circRNA reliability, since template switching events occur randomly during reverse transcription *in vitro*, a circRNA candidate observed in multiple samples or replicates is less likely to be generated from such an *in vitro* artifact. For the second most important factor in affecting circRNA reliability, full-length circular sequences can be reconstructed by Illumina short RNA-seq reads with the reverse overlap (RO) feature (Zheng et al. 2019) or identified by single-molecule long reads (Liu et al. 2021). With the support of RO-merged reads, the BSJ events are less probable to originate from template switching because both ends of the paired-end reads are sequencing from the same fragment and reversely overlapped with each other by alignment. Similarly, with the direct support of full-length circular sequences from single-molecule long reads, the nanopore-based full-length circRNAs are less likely to be artifacts derived from template switching.

In terms of circRNA biogenesis, we showed that the factors of both BSJ donor and acceptor splice sites at the same co-linear transcript isoforms was a good indicator for determining circRNA reliability. Regarding the transcripts generated during the transcriptional process, in addition to *cis*-backsplicing, an observed intragenic NCL junction may also arise from *trans*-splicing (Yu et al. 2014; Chen et al. 2015). Although the majority of detected NCL events may be derived from *cis*-backsplicing, previous studies based on comparisons of poly(A)-selected and poly(A)-depleted RNA-seq data suggested that a considerable percentage of detected NCL junctions were derived from *trans*-spliced RNAs (Chuang et al. 2016; Chuang et al. 2018). Actually, a number of NCL junctions detected in human cells were experimentally validated to be generated by *trans*-splicing rather than *cis*-backsplicing (Takahara et al. 2000; Flouriot et al. 2002; Wu et al. 2014; Yu et al. 2014). As *trans*-splicing occurs between two or more separate precursor mRNAs (Horiuchi and Aigaki 2006; Gingeras 2009), it is possible that an observed NCL junction with its two splice sites from different co-linear transcript isoforms is derived from a *trans-spl*icing event. We observed that the full-length circRNAs with the support of RO-merged short reads or single-molecule long reads, which were principally not generated from *trans*-splicing, were significantly enriched for the BSJs with donor and acceptor splice sites at the same co-linear transcript isoforms (Supplemental Fig. S3).

On the other hand, we showed that the factors related to RCS and RBPs binding to the flanking regions of BSJs were not good indicators for assessing circRNA reliability. Our observations reflect a previous notion that the existence of RCS is neither sufficient nor necessary for *cis*-backsplicing (Chen 2020). Although some circRNA cases were demonstrated to be facilitated by base-paring between RCSs at two flanking intronic sequences of the BSJs (Jeck et al. 2013; Zhang et al. 2014; Chuang et al. 2018), only a limited number of RCS-circRNA correlations were experimentally confirmed. Multiple RNA parings derived from different RCS pairs across flanking introns or within individual flanking introns can compete against each other and lead to the reduction of circRNA production (Ashwal-Fluss et al. 2014; Zhang et al. 2014; Kelly et al. 2015) or alternative *cis*-backsplicing (Zhang et al. 2016a; Zhang et al. 2020), further complicating the determination and validation of the RCS-circRNA correlations. In addition, while repetitive elements abundantly emerge in mammalian genomes, which contribute to RNA pairing, circRNAs observed in non-mammalian species (e.g., *Drosophila melanogaster* (Westholm et al. 2014) and *Oryza sativa* (Lu et al. 2015)) are often lacking the feature of RCS. For the associations between RBPs and circRNA formation, different RBPs may play opposite roles (up- or down-regulation) in regulating circRNA expression. Some RBPs, such as FUS (Errichelli et al. 2017) and ADARs (Shen et al. 2022), were even showed to mediate circRNA production in a bidirectional manner. Since different RBPs may have overlapping capacities for binding to RCSs, it requires further investigations to understand the joint (coordinate or competitive) effects of different RBPs on the regulations for each individual circRNA (Chen 2020; Yang et al. 2022).

This study systematically assessed the impacts of a dozen factors related to identification, conservation, biogenesis, and function on circRNA reliability on the basis of comparisons of mock-treated sample pairs from three different RNA treatment approaches. Of the examined factors, eight were shown to be the important indicators of circRNA reliability in all mock-treated sample pairs examined, regardless of the RNA treatment approaches. We assessed the relative influence of each individual factor on circRNA reliability and suggested that the most important factors in descending order were conservation level of circRNA, full-length circular sequences, supporting BSJ read count, both BSJ donor and acceptor splice sites at the same colinear transcript isoforms, supporting functional features, both BSJ donor and acceptor splice sites at the annotated exon boundaries, BSJs detected by multiple tools, and both BSJ donor and acceptor splice sites undergoing annotated AS events. All the circAtlas circRNA candidates and the corresponding factors examined were provided (Supplemental Table S3). Importantly, we found a remarkably positive correlation between number of supporting factors and the percentage of high-confidence circRNAs (i.e., RT-independent and RT-/non-RT-validated circRNAs) and a remarkably negative correlation between number of supporting factors and the percentage of circAtlas-specific circRNAs (Fig. 8). This revealed the additive effects of these factors on circRNA reliability. Particularly, more than 70% of circRNAs without any supporting factors were circAtlas-specific circRNAs, whereas such a percentage sharply declined to ≤3% when considering the circRNAs with at least five supporting factors (Fig. 8), supporting the effectiveness of these factors in determining circRNA reliability before performing experimental validations. To the best of our understanding, this study, for the first time, presents a useful guideline to systematically assess circRNA reliability with simultaneously accounting for varied types of factors, facilitating further functional investigation of this important class of non-canonical transcripts.

**Figure 8.**
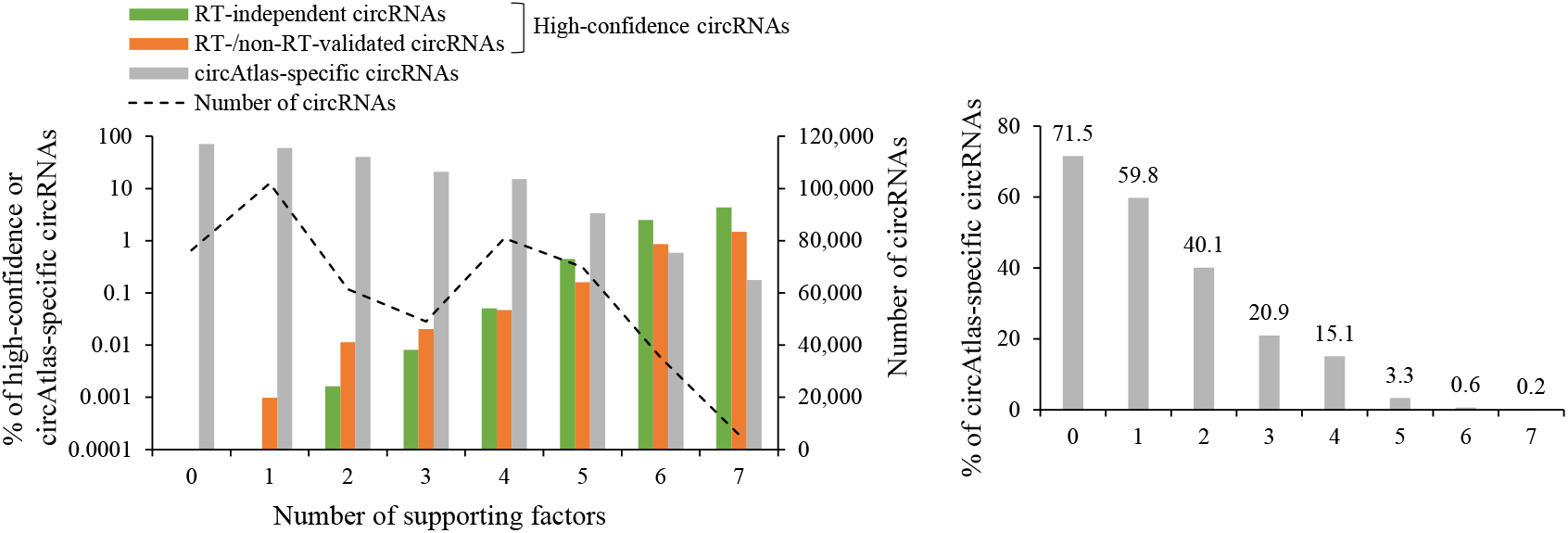
The correlation between number of supporting factors and the percentage of high-confidence circRNAs (RT-independent or RT-/non-RT-validated circRNAs) and circAtlas-specific circRNAs for the 480,471 circAtlas circRNA candidates. The right panel showed a clearer graph of the correlation between number of supporting factors and the percentage of circAtlas-specific circRNAs. The eight important factors illustrated in Fig. 6 except for the factor of “supporting BSJ read count” were considered, because this factor was dependent on the examined samples and the corresponding RNA-seq data. For the factors of “number of samples” and “number of supporting functional features”, three was used as a cutoff value. The detailed information can be found in Supplemental Table S3.

## Methods

### CircRNA candidates and RNA-seq data collection

As shown in Fig. 1A, human circRNA candidates examined (580,654 events; Supplemental Table S1) and the related information such as circRNA identification factors (e.g., circRNA detection tools and full-length circular sequences) and conservation factors (e.g., conservation of circRNAs across species, tissues, and samples) were downloaded from circAtlas 2.0 (Wu et al. 2020). CircAtlas circRNAs were identified by CIRI2 (Gao et al. 2015; Gao et al. 2018), find_circ (Memczak et al. 2013), circExplorer (Dong et al. 2019), or DCC (Cheng et al. 2016). The full-length circular sequences were reconstructed by CIRI-full (Zheng et al. 2019). All the analyses were based on the human reference genome (GRCh38) and the Ensembl annotation (version 100). The circRNA candidates that were potentially derived from alignment ambiguity (circRNA candidates had an alternative co-linear explanation or multiple matches against the human genome) were detected by NCLcomparator (Chen and Chuang 2019b) with default parameters and were not included in the subsequent analyses for accuracy (Fig. 1A). After that, 480,471 circAtlas circRNAs were retained. The three groups of mock-treated sample pairs (19 pairs) were extracted from publicly accessible databases (Table 1 and Supplemental Table S2). The downloaded mock-treated sample pairs should simultaneously satisfy the following criteria: (1) the number of the circRNA candidates detected in the mock samples should be greater than 600; and (2) more than one third of the circRNA candidates detected in the mock samples should be detected in the corresponding treated samples. For each RNA-seq dataset, duplicated RNA-seq reads were removed using FastUniq (Xu et al. 2012). The remaining RNA-seq reads were aligned against the human reference genome using STAR with default parameters (Dobin et al. 2013). For each circRNA (BSJ), we generated a pseudo sequence by concatenating the sequences flanking the BSJ (within −50 nucleotides of donor site to +50 nucleotides of acceptor site) and aligned all STAR-unmapped reads against the 100 bp BSJ-based pseudo sequence using BWA with default parameters (Li and Durbin 2009). A STAR-unmapped read was determined as a BSJ read if the read matched to ≥80% of the BSJ-based pseudo sequence and spanned the junction boundary by ≥10 bp on both sides of the BSJ.

### Treat/mock ratio calculation

For accuracy, only the circRNAs detected in the mock samples with the support of at least two BSJ reads were considered in the analysis for each mock-treated sample pair. The expression levels of circRNAs were calculated using the BSJ reads per million raw reads (RPM). The treat/mock ratios were defined as the ratio of the circRNA expression detected in the treated samples to that detected in the corresponding mock samples. The not-depleted and depleted circRNAs were defined as follows:

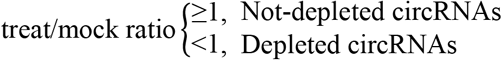

### Extractions of various factors

The factors of CIRI-full-identified full-length circular sequence, number of circRNA detection tools, conservation patterns at the individual (or sample), tissue, and specie levels (including tissue specificity index) were extracted from circAtlas 2.0 (Wu et al. 2020). The BSJ events with the direct support of full-length circular sequences from nanopore long RNA-seq reads were downloaded from Liu et al.’s study (Liu et al. 2021). The evolutionary rates were determined by phyloP (Pertea et al. 2011) or phastCons (Siepel et al. 2005) scores, which were downloaded from the UCSC Genome Browser at https://genome.ucsc..edu. For each BSJ, we considered the four regions around the BSJ: (1) within +1 to +10 nucleotides of the acceptor site; (2) within −10 to −1 nucleotides of the acceptor site; (3) within −10 to −1 nucleotides of the donor site; and (4) within +1 to +10 nucleotides of the donor site. The evolutionary rate of each region was measured by the average value of the phyloP (or phastCons) scores of the considered nucleotides (10 bp) within the region. The annotated exon boundaries and alternative splicing (AS) patterns were derived from the Ensembl annotation (version 100). The splice site strength of BSJs was estimated using MaxEntScan based on three scoring models (Maximum Entropy Model, First-order Markov Model, and Weight Matrix Model) (Yeo and Burge 2004). The numbers of reverse complementary sequences (RCSs) across the flanking sequences (RCS_across_; ±20,000 nucleotides of the BSJs) or within individual flanking sequences (RCS_swithin_) of the BSJs were calculated using CircMiMi (Chiang et al. 2022) with the command line: circmimi_tools check RCS --dist 20000 genome.fa circRNAs.tsv out.tsv. RBPs binding to the flanking regions (±1,000 nucleotides) of BSJs was determined according to CLIP-supported RBP binding sites downloaded from ENCORI (Li et al. 2014) at http://starbase.sysu.edu.cn/. miRNA binding sites across the BSJs were determined using miRanda 3.3a (Enright et al. 2003) with pairing score ≥ 155. RBP binding sites across the BSJs were determined using RBPmap (Paz et al. 2014) with high stringency level and conservation filter. The predicted miRNA (or RBP) binding sites should span the BSJ boundaries by ≥5 (or ≥2) bp on both sides of the BSJs, respectively. The seven types of evidence for coding potential of more than 320,000 human circRNAs (including circAtlas circRNAs) were extracted from TransCirc (Huang et al. 2021). The seven types of evidence included ribosome/polysome binding evidence, experimentally supported translation initiation site, internal ribosome entry site, predicted m6A modification site, circRNA-specific ORF, sequence composition score for the ORF, and mass spectrometry data-supported peptide across the BSJ. The translatable scores of circRNAs were downloaded from TransCirc.

### Extractions of high-confidence circRNAs and circAtlas-specific circRNAs

The high-confidence circRNAs examined here included RT-independent and RT-/non-RT-validated circRNAs (see text). The RT-independent circRNAs (1,478 events), which were supported by both Avian Myeloblastosis Virus (AMV)- and Moloney Murine Leukemia Virus (MMLV)-derived RNA-seq reads, were extracted from our previous study (Chuang et al. 2018). The RT-/non-RT-validated circRNAs (553 events), in which the NCL junctions (or their functions) were validated by RT-based experiments and at least one type of non-RT-based experiments in human tissues/cell lines, were extracted from CircR2disease (version 2.0) (Fan et al. 2021). The circAtlas-specific circRNA candidates were the BSJ events that were collected in circAtlas only rather than in eight other publicly-accessible circRNA databases (including CIRCpedia v2 (Dong et al. 2018), CircRic (Ruan et al. 2019), MiOncoCirc v2.0 (Vo et al. 2019), TSCD (Xia et al. 2017), circBase (Glazar et al. 2014), circRNADb (Chen et al. 2016), and exoRBase 2.0 (Lai et al. 2022). All the circRNA candidates deposited in CircR2disease or non-circAtlas circRNA databases were not considered if the NCL junction coordinates were not available or cannot be converted to the genomic coordinates on the GRCh38 assembly by liftOver (Hinrichs et al. 2006). For the NCL junctions with non-assigned strands, the junction coordinates were matched to the circRNA candidates in circAtlas. The analyzed circRNA candidates, the related factors, and the treat/mock ratios of the corresponding mock-treated sample pairs were deposited in Supplemental Table S3. All the coordinates in the table were one-based.

### Statistics analysis

For enrichment analyses, we compared not-depleted circRNAs with depleted ones in terms of various factors related to circRNA identification, conservation, biogenesis, and function. *P* values were determined using two-tailed Fisher’s exact test (FET) or Wilcoxon rank-sum test (WRST; greater and less) (see also the figure legends). *P* values were further FDR adjusted across 19 mock-treated sample pairs for each examined feature using Benjamini-Hochberg correction. To measure the relative effect of individual factor in determining circRNA reliability, we first employed a generalized linear model (GLM) with all the examined factors using the statsmodels package (0.13.2) (Seabold and Perktold 2010) in python and then performed the relative contributions to variability explained (RCVE) (Kvikstad et al. 2007) as follows:

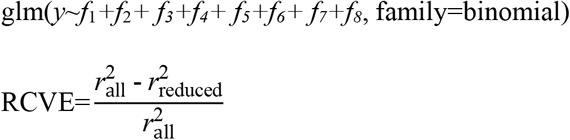

In GLM, *y* indicated whether a circRNA candidate was a not-depleted circRNA (i.e., not-depleted circRNA=1; depleted circRNA=0). *f*_1_~*f*_8_ represented the eight factors examined in the mode: supporting BSJ read count; whether BSJs were detected by multiple tools; whether BSJs can be reconstructed full-length circRNAs; number of samples observed the circRNAs; whether BSJs agreed to annotated exon boundaries; whether BSJs had both donor and acceptor splice sites matching annotated exon boundaries from the same co-linear transcript isoforms; whether BSJs were subject to AS; and number of supporting functional features, respectively. In the RCVE formula, 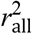 and 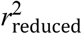 represented the *r*^2^ values calculated by GLM including all the eight factors and excluding the factor of interest, respectively. The RCVE values range from 0 to 1, with a higher RCVE value indicating a more important contribution of the examined factor to the model. The detailed results of the statistical significance tests for each examined factor were deposited in Supplemental Table S4.

## Data availability

The data underlying this study are available in the text and in Supplemental Tables 1-4. Supplemental Tables 1-4 and all the codes used to generate the results are available at GitHub (https://github.com/TreesLab/circRNA_features).

## Competing interest statement

The authors declare no competing interests.

## Acknowledgements

This work was supported by Genomics Research Center, Academia Sinica, Taiwan to T.-J.C. and the National Science and Technology Council, Taiwan (MOST 111-2311-B-001-021) to T.-J.C. We thank the related organizations for generating and providing the RNA-seq data used in this study. We also thank other members of Trees-Juen Chuang lab for helpful discussions.

## Notes

### Competing Interest Statement

The authors have declared no competing interest.

